# Selective culture enrichment and sequencing of faeces to enhance detection of antimicrobial resistance genes

**DOI:** 10.1101/560292

**Authors:** Leon Peto, Nicola J. Fawcett, Derrick W. Crook, Tim E.A. Peto, Martin J. Llewelyn, A. Sarah Walker

**Affiliations:** National Institute for Health Research (NIHR) Health Protection Research Unit on Healthcare Associated Infections and Antimicrobial Resistance, Oxford, UK; Nuffield Department of Medicine, University of Oxford, UK; National Infection Service, Public Health England, Colindale, UK; Department of Global Health and Infection, Brighton and Sussex Medical School, Falmer, UK; Department of Microbiology and Infection, Brighton and Sussex University Hospitals NHS Trust, Brighton, UK

## Abstract

Metagenomic sequencing of faecal DNA can usefully characterise an individual’s intestinal resistome but is limited by its inability to detect important pathogens that may be present at low abundance, such as carbapenemase or extended-spectrum beta-lactamase producing *Enterobacteriaceae*. Here we aimed to develop a hybrid protocol to improve detection of resistance genes in *Enterobacteriaceae* by using a short period of culture enrichment prior to sequencing of DNA extracted directly from the enriched sample. Volunteer faeces were spiked with carbapenemase-producing *Enterobacteriaceae* and incubated in selective broth culture for 6 hours before sequencing. Different DNA extraction methods were compared, including a plasmid extraction protocol to increase the detection of plasmid-associated resistance genes. Although enrichment prior to sequencing increased the detection of carbapenemase genes, the differing growth characteristics of the spike organisms precluded accurate quantification of their concentration prior to culture. Plasmid extraction protocols increased detection of resistance genes present on plasmids, but the effects were heterogeneous and dependent on plasmid size. Our results demonstrate methods of improving the limit of detection of selected resistance mechanisms in a faecal resistome assay, but they also highlight the difficulties in using these techniques for accurate quantification and should inform future efforts to achieve this goal.

## INTRODUCTION

The intestinal microbiome is an important reservoir for antimicrobial resistance (AMR) genes^1^. Accurate detection and quantification of AMR genes within an individual’s gut microbiome (their gut ‘resistome’) would allow the impact of different types of antibiotic exposures to be evaluated in both research and clinical practice, and could guide infection control interventions to reduce AMR. It would be particularly helpful for antibiotic stewardship, as current policies often focus on the use of drugs with a ‘narrow’ rather than ‘broad’ clinical spectrum, despite the fact that this may poorly reflect their impact on the gut microbiome and resistome^2^.

As AMR increases there is a growing need for standardised methods of resistome assessment that would allow meaningful comparisons between settings. However, there is currently no single method that can fully characterise the gut resistome. Quantitative culture requires further genotyping to define resistance elements and can only practically be used for a small number of species present in faeces^3^, and targeted molecular approaches, such as PCR, cannot simultaneously detect the dozens of AMR genes and mutations that may be present in a sample from among thousands of known variants^4^. Because of these shortcomings, metagenomic sequencing performed directly from extracted faecal DNA is increasingly being used to characterise the gut resistome, as it produces millions of short DNA reads that can be matched to a catalogue of thousands of AMR genes, theoretically producing a representative, quantitative description of AMR genes in a sample^1,5,6^.

However, a major limitation of direct sequencing is the inability to detect clinically important resistance genes present at low abundance^7^. In particular, the *Enterobacteriaceae*, which include major drug resistant human pathogens such as *Escherichia coli* and *Klebsiella* species, may be present in faeces at levels too low for AMR gene detection by direct sequencing. In one recent study, metagenomic sequencing detected an extended spectrum beta-lactamase (ESBL) in just 12 of 26 (46%) culture positive stool samples, missing all 9 samples with an abundance of <10^6^ CFU/g^8^. Increasing the depth of sequencing to detect scarce organisms seems appealing but is not currently feasible. For example, it would not be unusual for a resistant *E. coli* to be present at an abundance of 10^3^/g in faeces with a total organism count of 10^11^/g^8,9^, so using short read sequencing to detect a 1kb AMR gene present in this organism would require hundreds of billions of reads, costing many hundreds of thousands of dollars. We therefore aimed to assess in proof-of-principle experiments whether a short period of selective culture enrichment of faecal samples before sequencing could improve the detection and quantification of AMR in *Enterobacteriaceae* by raising their abundance above the threshold of detection.

## MATERIALS AND METHODS

### Specimens

The Antibiotic Resistance in the Microbiome – Oxford (ARMORD) Study gathers faecal samples from patients and healthy volunteers to study antimicrobial resistance with approval from Leicester Research Ethics Committee (reference 15/EM/0270). Stool from two ARMORD participants was used for all experiments other than the culture calibration experiment. Participant A was a hospital inpatient with exposure to a third-generation cephalosporin antibiotic 48 hours previously, participant B was a healthy volunteer with no recent antibiotic exposure. Samples were frozen at −80°C within 2 hours of collection and were culture-negative for ESBL producing *Enterobacteriaceae*. The culture calibration experiment used a discard faecal sample from the clinical microbiology laboratory at the Oxford University Hospitals NHS Foundation Trust.

### Quantitative culture

Modified track plating was used to quantify *Enterobacteriaceae* in faecal samples^10^. Samples were serially diluted 10-fold in peptone water with pipette mixing at each step. Culture of 10ul from dilutions 1:10^-1^ - 1:10^-8^ was performed on CHROMagar Orientation (BD) and ESBL Brilliance agar (Oxoid). Colonies were counted after incubation at 37°C for 16 hours in aerobic conditions.

### Experimental bacterial strains

Three carbapenemase-producing *Enterobacteriaceae* (CPE) strains were used for the spiking experiments, grown on CRE Brilliance agar (Fisher Scientific) at 37°C in aerobic conditions.

1. *E. coli* H17. Contains NDM-1 beta-lactamase on unresolved plasmid, isolated from a hospital outbreak in Nepal (SAMN02885348)^11^.
2. *Klebsiella pneumoniae* PMK3. Contains NDM-1 and CTX-M-15 beta-lactamases on a 305kb plasmid, present at an estimated 1.8 copies per chromosome. Isolated from a hospital outbreak Nepal (SAMN02885389)^11^.
3. *Enterobacter cloacae* CAV1668. Contains KPC-2 and TEM-1 beta-lactamases on a 43kb plasmid, present at an estimated 1.9 copies per chromosome. Isolated from a hospital outbreak in Virginia, USA (SAMN03733827)^12^.

### Spiking

Overnight growth of bacteria was used to make a suspension of 5 MacFarland in nutrient broth with 10% glycerol (NBG) (Oxoid). Serial 10-fold dilutions were frozen at −80°C with culture from thawed aliquots used to define concentrations. Stool samples were vortexed with molecular water (Fisher Scientific) on ice at a ratio of 1:2 to create a pipettable suspension, 300mg of which was used per set of experimental conditions. Samples were spiked to a final concentration of 5×10^2^, 5×10^3^, 5×10^4^, 5×10^5^, 5×10^6^, or 5×10^7^ CFU/g, using 5-50ul of spiking solution, or with 50ul NBG alone for unspiked controls. In addition, to allow quantification after sequencing, a standard spike of 10^7^ CFU of *Staphylococcus aureus* was added to all samples except negative controls immediately prior to DNA extraction. MRSA252 (SAMEA1705935) was used, which was grown on blood agar (Fisher Scientific) at 37°C in aerobic conditions.

### Enrichment

Spiked samples were added to 20ml Mueller-Hinton broth (Sigma-Aldrich) containing 20mg/L vancomycin, 20mg/L metronidazole, and for later spiking experiments, 1mg/L cefpodoxime (all antibiotics Sigma-Aldrich). These were incubated at 37°C and shaken at 190rpm for 6 hours, then immediately centrifuged at 3200g for 10 min at 4°C to pellet bacteria. Supernatant was discarded and the pellet resuspended in 1ml molecular water.

### DNA extraction

Two DNA extraction protocols were used, which were compared in an initial extraction experiment. The subsequent vancomycin and metronidazole enrichment experiment used the Fast DNA Stool Mini Kit, and the cefpodoxime, vancomycin and metronidazole enrichment experiment used the QuickLyse Miniprep kit.

1. QuickLyse Miniprep Kit (Qiagen). Samples were centrifuged at 100g for 5 minutes to remove large particles, and DNA was then extracted from the supernatant according to the manufacturer’s instructions. Briefly, bacterial cells were pelleted by centrifugation at 17,000g and resuspended in 400ul ice-cold complete lysis solution by vortexing for 30s. After 3 minutes at room temperature the lysate was transferred to a QuickLyse spin column and centrifuged at 17,000g for 60s. The spin column was washed with 400ul QuickLyse wash buffer and centrifuged at 17,000g for 60s. After discarding the flow-through, the columns were dried by centrifuging at 17,000g for 60s. DNA was eluted from the spin column in 50 ul QuickLyse elution buffer by spinning at 17,000g for 60s.
2. QIAamp Fast DNA Stool Mini Kit (Qiagen) plus mechanical lysis with bead beating. Samples were added to 1ml Stool Transport and Recovery buffer (Roche) in a 2ml Lysing Matrix E tube (MP Biomedicals), followed by bead beating for 2 x 40s with a FastPrep-24 5G instrument (MP Biomedicals) and incubation at 95°C for 5 minutes. Samples were then centrifuged at 1,000g for 60s. Supernatant was transferred to 1ml InhibitEX buffer, after which the manufacturer’s instructions were followed for bacterial DNA extraction. Briefly, this involved mixing with 15ul proteinase K, 400*µ*l AL lysis buffer and 400ul 100% ethanol. This was transferred to a QIAamp spin column and washed with AW1 and AW2 buffers. DNA was then eluted in molecular water.

### DNA library preparation and sequencing

Extracted DNA was purified using Ampure XP beads (Agilent). 50ul DNA was vortexed with 90ul Ampure XP beads and washed twice with 80% ethanol before elution in 25ul molecular water. DNA was normalised to 0.2ng/ul and libraries prepared from 1ug DNA using the Nextera XT Library preparation kit (Illumina) according to the manufacturer’s instructions. Normalisation and QC of libraries was performed using a Qubit 2.0 fluorometer (Thermo Fisher) and 2200 Tapestation (Agilent). Paired-end 300bp sequencing was performed on an Illumina MiSeq, with 10–16 samples multiplexed per sequencing run. Median sequencing depth ranged from 500,000 to 1.4 million 300bp paired reads.

### Bioinformatics

Human DNA reads (representing <4% of sequences in all samples) were identified using the Kraken taxonomic classifier and removed prior to analysis^13^. Quality control metrics were assessed with FastQC^14^ and BBduk was used to remove Illumina adapters and quality-trim reads, using a phred-score cut-off of 10^15^. MetaPhlAn2 was used for taxonomic classification^16^, except for normalisation to *S. aureus*, where Kraken was used to classify reads to the relevant species. BBMap was used to map reads to carbapenemase genes, *gyrA* and plasmids^15^. Plasmid copy number was estimated by mapping Illumina reads from sequenced isolates to a closely related closed reference genome using BWA-MEM^17^. For PMK3, the PMK1 reference was used (CP008929-CP008933), and for CAV1668, the CAV1668 reference was used (CP011582-CP011584). SAMtools was used to filter out supplementary alignments and calculate depth of coverage^18^.

### Normalisation to *S. aureus*

To allow quantification of CPE spike organisms after enrichment, a standard inoculum of 10^7^ CFU *S. aureus* was added to all samples after enrichment and immediately prior to DNA extraction. Detection of DNA reads from different organisms depends on several factors such as genome size, efficiency of DNA extraction, efficiency of sequencing and efficiency of bioinformatic classification. These can be combined into a single measure of relative detection efficiency (RDE) compared to *S. aureus*, which is established using unenriched samples in which the concentration of the spike organism and *S. aureus* are both known. For each CPE spike organism, the mean RDE was determined from 4 unenriched samples spiked with 5×10^7^CFU/g CPE and 10^7^ CFU *S. aureus* (shown in **Fig 4**), using the formula:

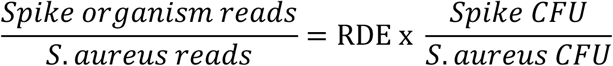

The RDE was then used to estimate the number of CFUs present in samples after enrichment.

## RESULTS

### Comparison of DNA extraction methods

Preliminary experiments were performed to identify a DNA extraction protocol that would efficiently extract resistance genes present in *Enterobacteriaceae*. As acquired resistance genes in *Enterobacteriaceae* are typically present on plasmids, we hypothesised that a plasmid extraction protocol would increase detection of these genes. The QuickLyse Miniprep Kit (Qiagen), a spin column-based plasmid extraction kit, was compared to a standard faecal DNA extraction protocol, the QIAamp Fast DNA Stool Mini Kit (Qiagen) modified to include mechanical lysis with bead beating. One hundred milligram aliquots from two different volunteer faecal sample were spiked with one of two *Enterobacteriaceae* carrying plasmid-associated beta-lactamases (*K. pneumoniae* with NDM-1 and CTX-M-15, or *E. cloacae* with KPC-2 and TEM-1). Spike organisms were added at a concentration of 10^8^ CFU/g and negative controls with no spike were run in parallel. Following DNA extraction using the two methods, sequencing was performed on Illumina MiSeq. Reads mapping to the plasmid-associated beta-lactamases were increased using the QuickLyse kit, to a mean of 15.7 Read per Kb per million (RPKM) (95% CI 6.2 – 25.1) compared to 4.4 RPKM (95% CI 2.4 – 6.4, t-test for difference p=0.03) using the Fast DNA Stool kit (**Supplementary Fig**).

### Selective enrichment with vancomycin and metronidazole

Before assessing a protocol of enrichment-culture and direct sequencing, we first had to determine the optimal duration of culture. We therefore performed a calibration culture experiment in which a 100mg of a faecal sample was incubated in Mueller-Hinton broth containing metronidazole (20mg/L) and vancomycin (20mg/L), repeated in duplicate. These antibiotics were added to suppress the growth of anaerobic and gram-positive organisms but allow growth of *Proteobacteria*, which would primarily consist of *Enterobacteriaceae*. Quantitative culture was performed at intervals over 24 hours, and demonstrated that *Enterobacteriaceae* were in log-phase growth up to approximately 8 hours of incubation (**Fig 1**). Six hours was selected as the optimal incubation period, as this provided approximately 10^4^-fold enrichment while preserving the ratios of the different *E. coli* species present at baseline.

**Figure 1.**
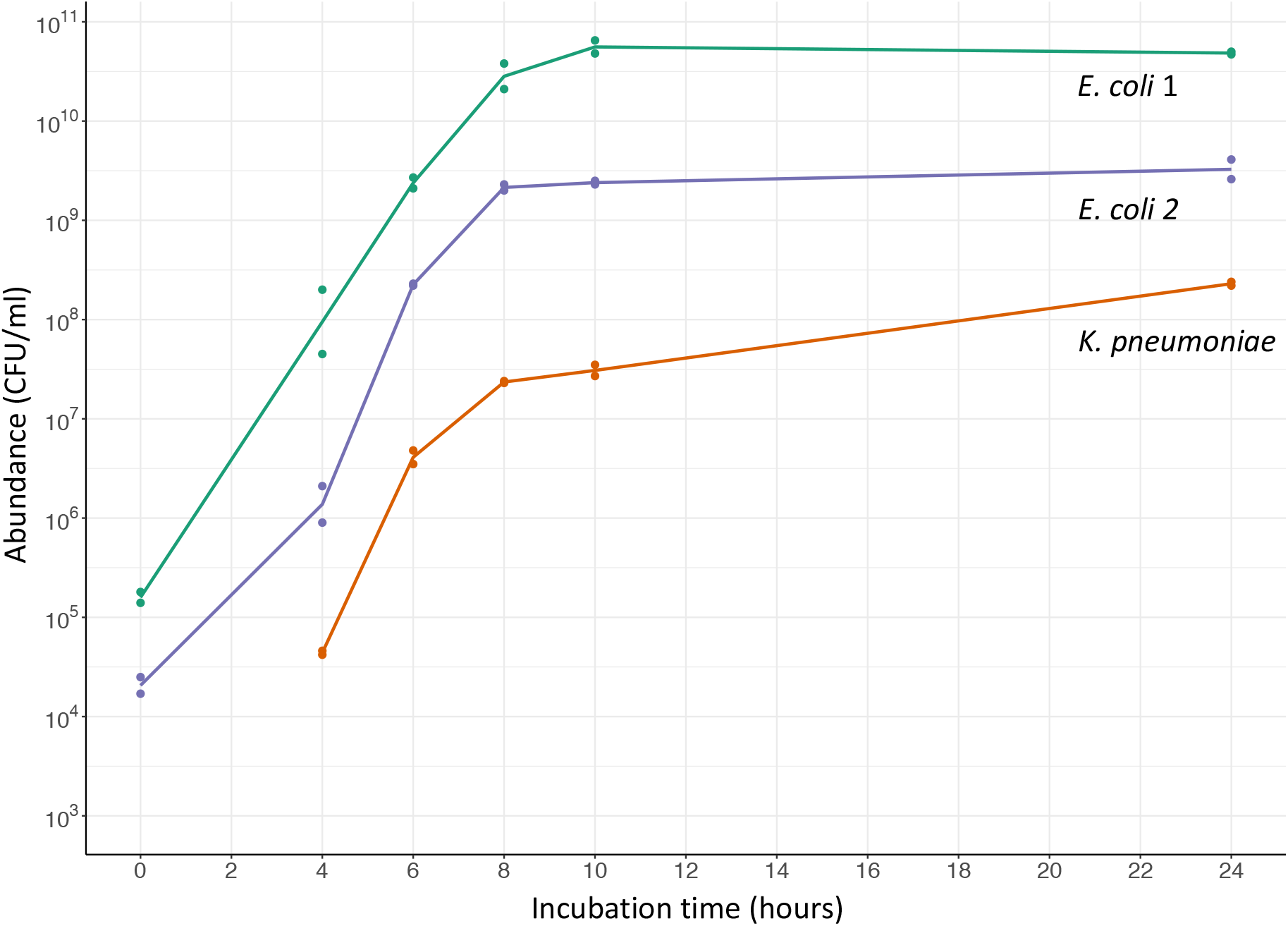
Growth of *Enterobacteriaceae* during incubation of a single faecal sample in broth containing vancomycin and metronidazole. Two distinct morphotypes of *E. coli* were present. No *K. pneumoniae* were isolated at 0 hours, implying presence below the limit of detection of 10^3^ CFU/g. Lines join geometric means of 2 replicates.

We then assessed the effect of this selective enrichment protocol on detection of AMR genes present in *Enterobacteriaceae* by sequencing. One hundred milligram aliquots from a single faecal sample were spiked with *E. coli* containing NDM-1 at concentrations of 0 – 5×10^5^ CFU/g, repeated in triplicate. These had 6 hours of enrichment culture in broth containing metronidazole and vancomycin, followed by DNA extraction with the Fast DNA Stool kit and sequencing on Illumina MiSeq. This showed that at all spiking concentrations tested, the spike organism was barely detectable above endogenous *E. coli* present in the sample, with a relative abundance of *E. coli* of around 3% in all unenriched samples, and around 98% in all enriched samples (**Fig 2**). The NDM-1 gene was scarcely measurable in any samples, being detected in just a single read in 3 enriched samples: 2 of 3 spiked at 5×10^5^ CFU/g, and 1 of 3 spiked at 5×10^4^ CFU/g, in each case corresponding to an abundance of <1 RPKM (starred samples in **Fig 2**).

**Figure 2.**
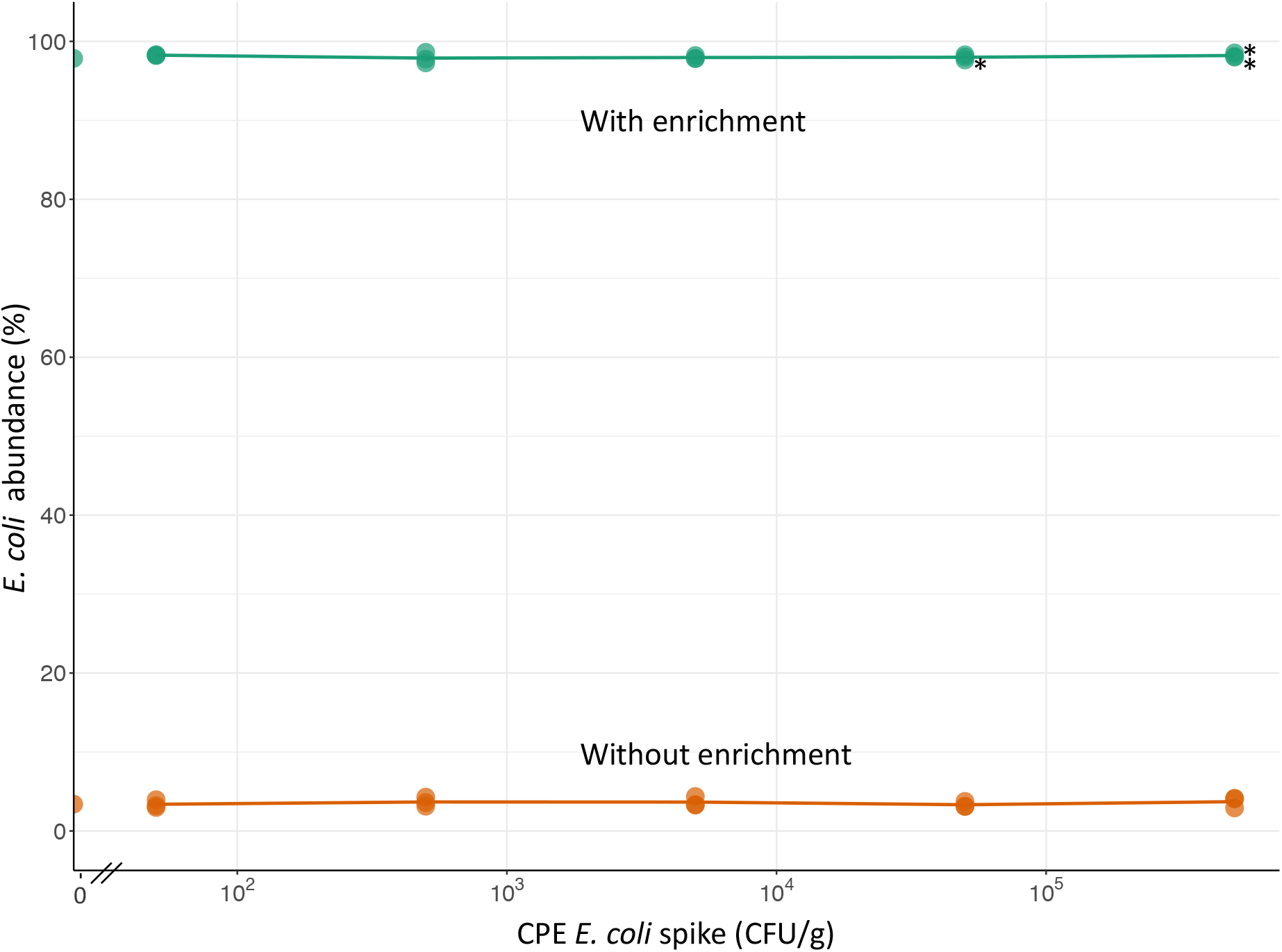
*E. coli* abundance in aliquots of a single faecal sample spiked with NDM-1 *E. coli*, with or without 6 hours of enrichment in broth containing vancomycin and metronidazole. Note that the sample contained endogenous non-CPE *E. coli*. Stars indicate the three samples in which a single DNA read mapped to the NDM-1 gene, with no reads mapping to the gene in any other samples. Lines join arithmetic means of 3 replicates, unspiked samples are shown on the y-axis.

### Selective enrichment with vancomycin, metronidazole and cefpodoxime

In order to detect subpopulations of *Enterobacteriaceae* resistant to third-generation cephalosporins, a revised protocol added cefpodoxime (1mg/L) to suppress growth of susceptible *Enterobacteriaceae* and also used the QuickLyse plasmid extraction method above. To establish its performance, another spiking experiment was performed, this time using faecal samples from two volunteers. One hundred milligram aliquots from the samples were spiked with one of two different carbapenemase-producing *Enterobacteriaceae* (NDM-1 *K. pneumoniae* or KPC-2 *E. cloacae*) at spikes ranging from 0 – 5×10^7^ CFU/g. Unenriched controls were run in parallel and all conditions were repeated in duplicate. Immediately before sequencing, 10^7^ CFU *S. aureus* was added to all samples as a standard to allow quantification of the spike CPE.

Using this revised protocol, CPE spikes above 5×10^5^ CFU/g led to a relative abundance of ≥90% of the spike organism after enrichment (**Fig 3**). At lower spiking concentrations, there was a marked difference between *K. pneumoniae* and *E. cloacae*. At an initial spike of 5×10^2^ CFU/g, *K*. *pneumoniae* had a mean abundance of 58% compared to 3% for *E. cloacae*. In all unenriched samples the spike organism was present at an abundance of <10%.

**Figure 3.**
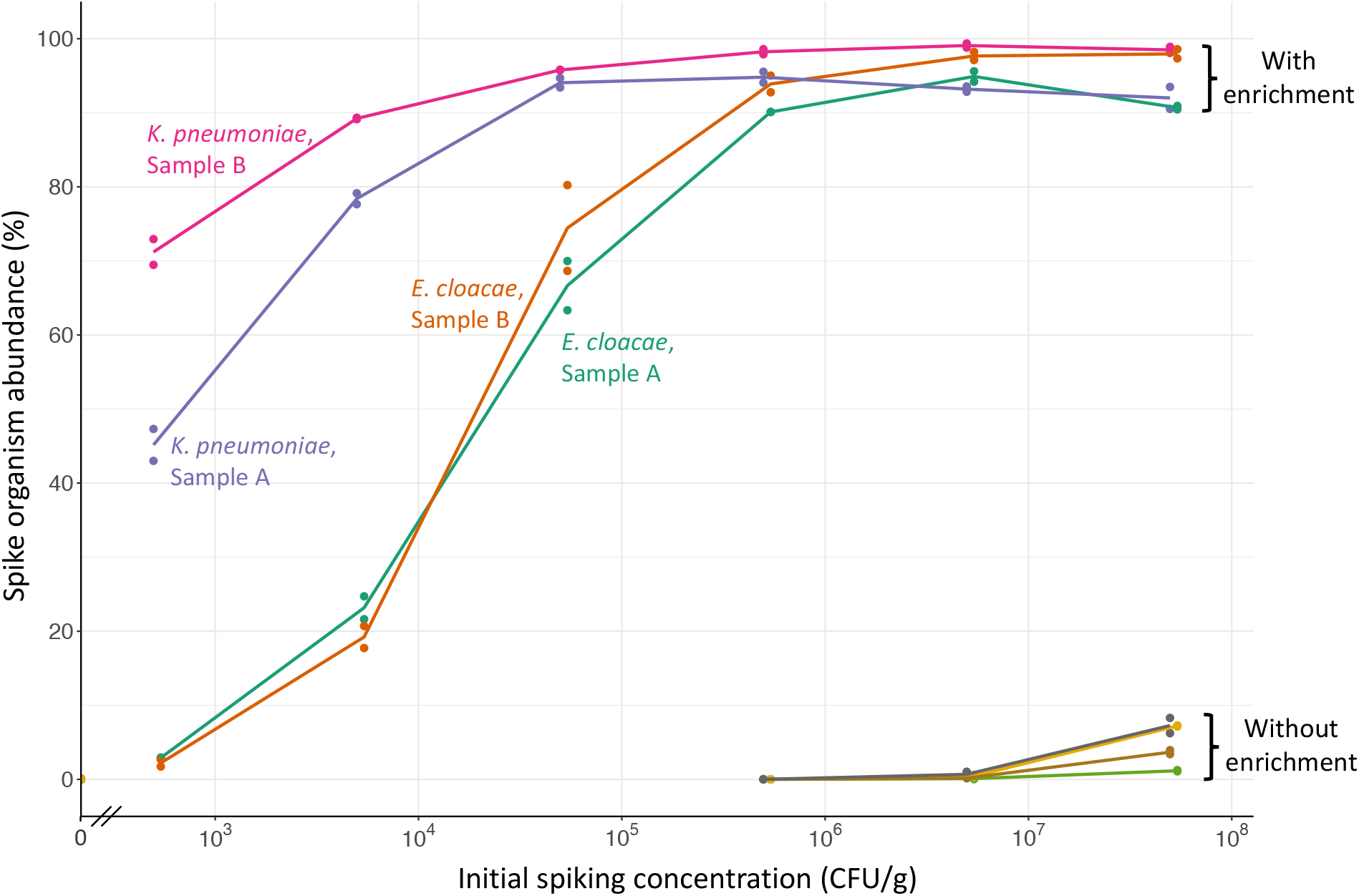
Initial CPE spike versus measured abundance of spike organism, with or without enrichment in broth containing cefpodoxime, vancomycin, and metronidazole. Two spike CPE (*K. pneumoniae* & *E. cloacae*) and two volunteer samples (A & B) were used. Lines join arithmetic means of 2 replicates, unspiked samples are shown on the y-axis.

Our objective was to measure the absolute, as well as relative, abundance of organisms present in samples, which requires comparison to a known standard. We estimated the abundance of spike organisms after enrichment by comparing reads assigned to the spike to those assigned to the *S. aureus* standard (see Methods). Abundance estimation in the enriched samples showed that the number of spike organisms increased with higher initial spiking concentration even after the relative abundance had plateaued (**Fig 4**). Comparing these estimates to measurement by quantitative culture showed reasonable correlation at abundances below 10^9^ CFU, corresponding to initial spikes <5×10^5^ CFU/g, with all estimates within 1 log of the culture value (**Fig 5**). However, at higher abundances, where spike organisms outnumbered the standard more than 300-fold, sequencing consistently underestimated culture values.

**Figure 4.**
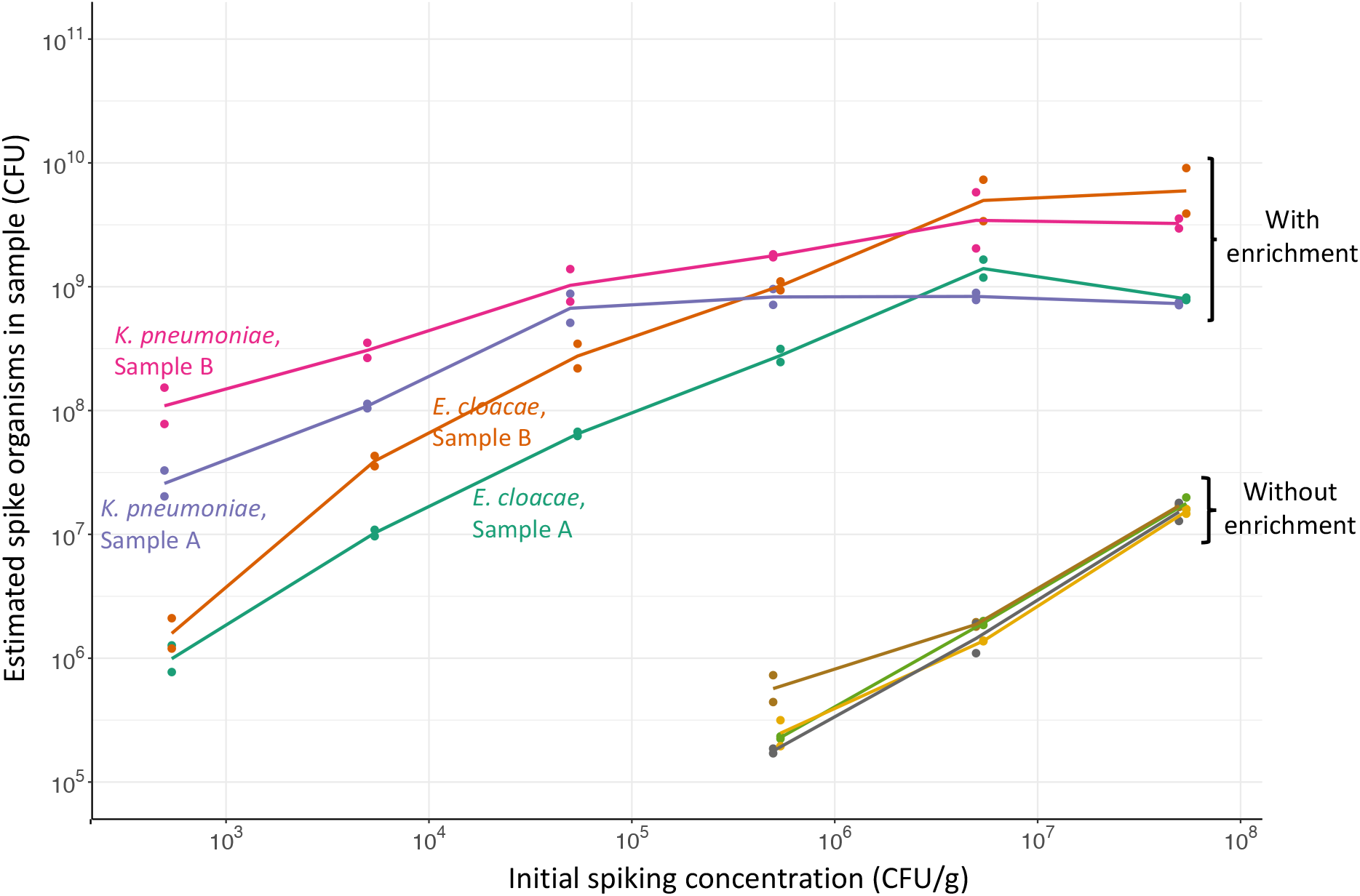
Initial CPE spike versus final number of CPE spike organisms estimated from sequence data by normalisation to *S. aureus*. With and without enrichment in broth containing cefpodoxime, vancomycin, and metronidazole. Two spike organisms (*K. pneumoniae* & *E. cloacae*) and two volunteer samples (A & B) were used. Lines join geometric means of 2 replicates.

**Figure 5.**
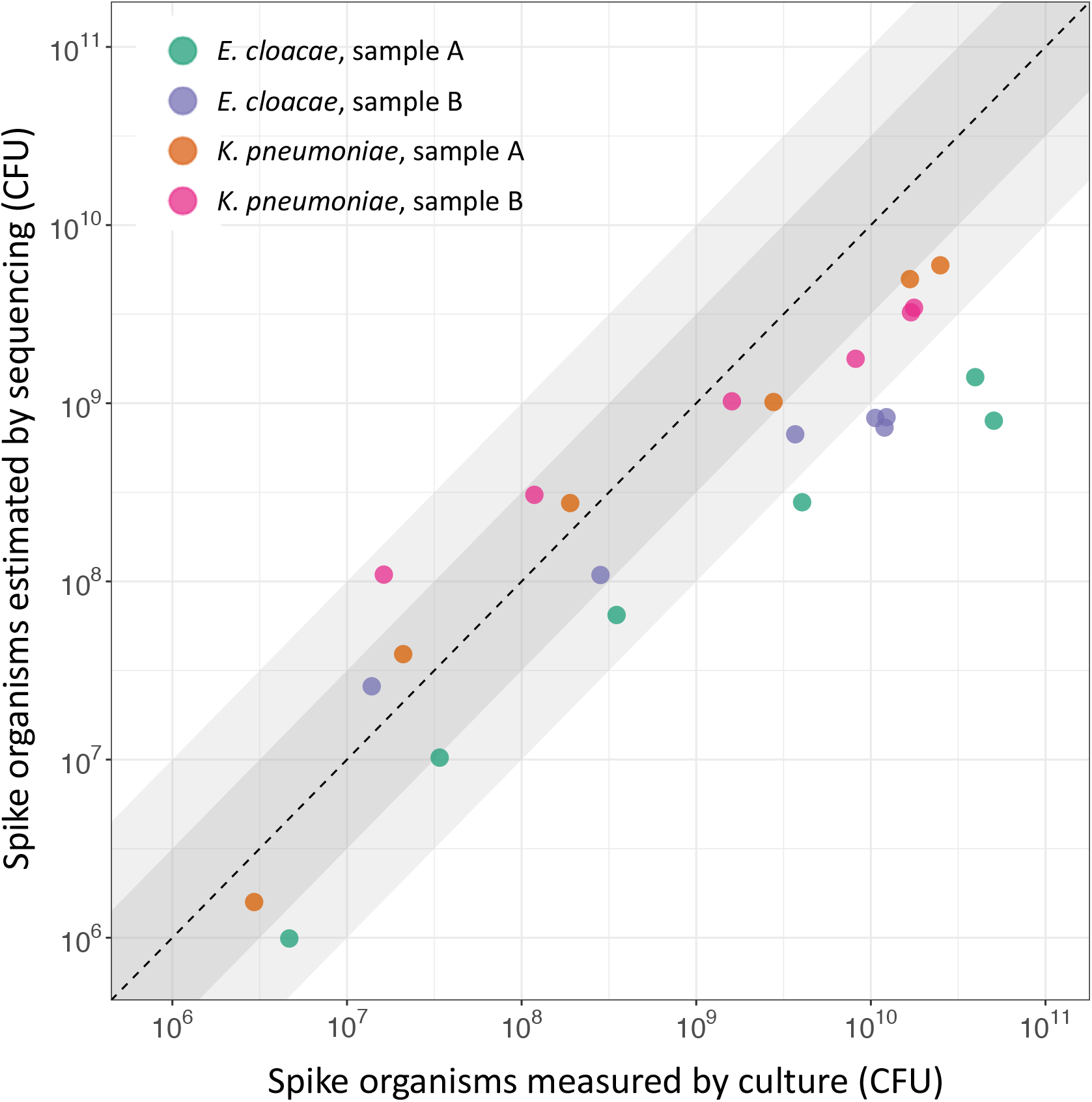
Number of spike organisms estimated by sequencing compared to measurement by culture. Samples enriched in broth containing cefpodoxime, vancomycin, and metronidazole. Two spike organisms (*K. pneumoniae* & *E. cloacae*) and two volunteer samples (A & B) used. Points represent geometric means of 2 replicates, shaded areas are within 0.5 and 1.0 log_10_ of the line of identity.

Detection of the carbapenemase genes differed markedly between the two spike organisms (solid lines in **Fig 6**). The KPC-2 gene present in *E. cloacae* was detected at a mean 12 RPKM (range 5-17) at the lowest spike of 5×10^2^ CFU/g, implying a limit of detection of approximately 50 CFU/g per million reads. In contrast, the NDM-1 gene present on *K. pneumoniae* was detected at around 100 RPKM even at the lowest initial spike and hardly changed with higher spiking concentrations, in keeping with the higher abundance of *K. pneumoniae* noted above. This prevented estimation of the limit of detection of NDM-1 in this context.

**Figure 6.**
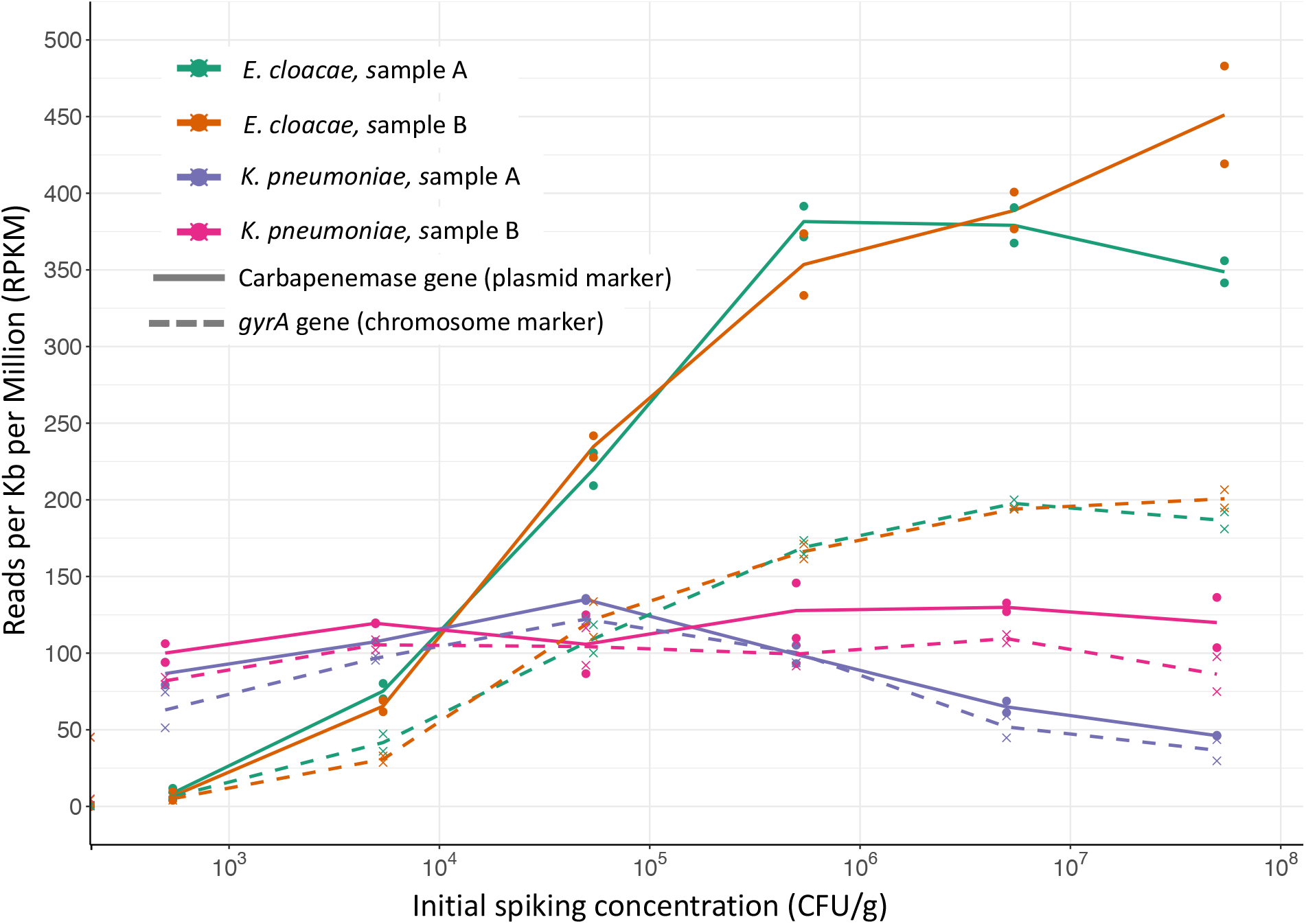
Initial CPE spike versus reads mapping to carbapenemase genes (solid lines, NDM-1 in *K. pneumoniae*, KPC-2 in *E. cloacae*), or gyrA (dashed lines). Samples enriched in broth containing cefpodoxime, vancomycin, and metronidazole. Two spike organisms (*K. pneumoniae* & *E. cloacae*) and two volunteer samples (A & B) were used. Lines join arithmetic means of 2 replicates, unspiked samples are shown on the y-axis.

At the highest spiking concentrations, the number of reads mapping to the carbapenemase was around 4 times higher in *E. cloacae* than *K. pneumoniae*, even though both organisms were present at >90% abundance. This difference did not appear to relate to the plasmid copy number, as in sequenced isolates the carbapenemase genes are present at similar copy numbers of 1.8 – 1.9. To explore the possibility that this difference related to efficiency of plasmid extraction, we compared the number of reads mapping to the plasmid-associated carbapenemase to those mapping to the chromosomal *gyrA* gene (dashed lines in **Fig 6**). In *E. cloacae*, DNA from the 43kb KPC plasmid was extracted more efficiently than the chromosomal *gyrA* gene, leading to greater detection of KPC-2 vs gyrA. In contrast, in *K. pneumoniae* DNA from the 305kb NDM-1 plasmid and *gyrA* was extracted at similar efficiencies.

As well as the NDM-1 plasmid, the *K. pneumoniae* spike also contains DNA mapping to two small plasmids, pEC34A (3.8kb) and pRGRH1815 (7.1kb), although the structure of these in the spike organism could not be resolved with available sequence data^19^. This DNA was extracted at a very high efficiency in enriched samples, and the extraction efficiency relative to chromosomal DNA increased with higher spiking concentrations (**Fig 7**). At the highest spikes this small plasmid DNA made up around a quarter of all reads, causing a relative decrease in other *K. pneumoniae* DNA.

**Figure 7.**
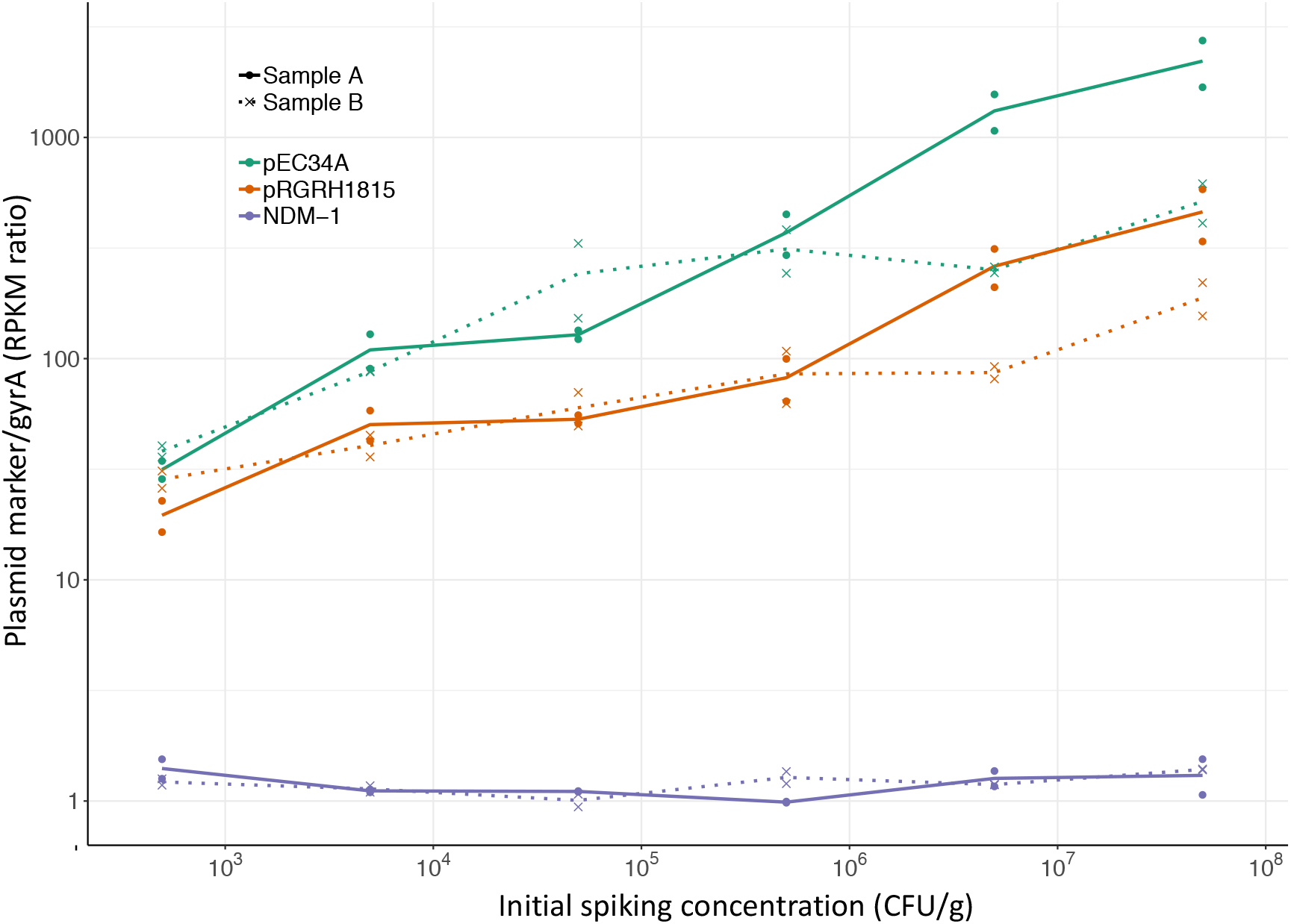
Initial *K. pneumoniae* spike versus ratio of plasmid-marker to chromosomal gyrA reads. Samples enriched in broth containing cefpodoxime, vancomycin, and metronidazole. Two volunteer samples (A & B) were used. Lines join geometric means of 2 replicates.

## DISCUSSION

Tools to rapidly and reliably assay an individual’s faeces for carried antibiotic resistance would allow personalised assessment of the risks of antibiotic exposure, target infection control activity and aid research to objectively compare the impact of different treatment strategies on resistance. Standardised measures of this selection pressure across a wide range of antibiotics and clinical settings could greatly inform prescribing choices^2^. While microbiome measurements have become relatively standardised, using either 16S PCR or shotgun metagenomic sequencing, there is no standardised resistome assay, because all current methods suffer from serious limitations. Quantitative culture requires further genotyping to define resistance elements, and cannot detect resistance genes in non-target organisms. PCR can only target a limited number of resistance mechanisms in parallel, and although direct sequencing can detect a large panel of AMR genes, it cannot detect organisms present at low abundance^8^.

Here we have assessed a novel resistome assay combining a short period of selective culture with sequencing directly from the enriched sample. We chose culture conditions that would amplify a resistant subpopulation of *Enterobacteriaceae* of particular clinical importance, specifically those resistant to third generation cephalosporins, with the aim of bringing their resistance genes above the threshold of detection. The relative abundance of resistance genes within the faecal microbiome is expected to be associated with antibiotic exposure^20^, risk of invasive infection^21^ and transmission^22^ so it is important that enrichment of resistant organisms is not at the expense of quantification. Therefore, after initial experiments to define the growth of *Enterobacteriaceae* in selective broth culture, we chose an incubation period that would produce a 10^4^-fold increase whilst maintaining log-phase growth to allow quantification.

Performance of the enrichment-sequencing protocol was assessed by spiking faecal samples with known concentrations of CPE. While there were minimal differences between samples from different individuals, there were large differences between the two spike organisms. Consistency between the faecal samples, and between replicates, implies that this relates to intrinsic growth characteristics of the strains used in this experiment. Our finding of such large variation in growth between different strains of *Enterobacteriaceae* precludes accurate estimation of their starting concentration unless their growth characteristics are known, undermining one of the aims of the assay. In fact, relatively small differences in growth rates could produce such differences after 6 hours. For example, a doubling time of 18 minutes versus 22 minutes would lead to a 12-fold difference in abundance at 6 hours. As well as intrinsic differences between bacterial strains, variation in growth rate can also be influenced by total bacterial concentration^23^ and the faecal environment^24^.

To measure the absolute abundance of organisms in the sample it is necessary to use some form of standard. Although our use of a *S. aureus* standard was able to estimate the number of post-enrichment spike organisms at low abundances, it correlated poorly with quantitative culture at higher abundances. This is likely to be related to small amounts of contaminating DNA assigned to the standard, as even in negative controls with no faecal sample a small number of reads were assigned to *S. aureus*. This could possibly be improved by using an alternative standard with less homology to any possible contaminants or organisms present in the sample.

Another novel aspect of this protocol was the use of plasmid DNA extraction, which increased the detection of plasmid-mediated resistance in *Enterobacteriaceae* in preliminary spiking experiments. To our knowledge this is the first study to compare plasmid DNA extraction to standard DNA extraction for resistome assessment in humans. One study assessing AMR genes in environmental metagenomes found that plasmid extraction could increase detection of some genes 37 fold^25^, but in our study the effect on the genes of interest was much more modest, with increases of only 3-4 fold. More problematically, plasmid DNA extraction produced unwanted artefacts because of differential extraction efficiency depending on plasmid size. This was noticeable when comparing the 305kb and 43kb carbapenemase containing plasmids, but was much more marked in relation to reads mapping to small 4-7kb plasmids, which were extracted at extremely high efficiency compared to other DNA.

This study has several limitations. It was only possible to assess a limited number of conditions, and the quantitative findings of this study may have been different if using different spike organisms, culture conditions or plasmid extraction methods, although it seems unlikely that the qualitative conclusions would differ. The use of a plasmid extraction kit in some experiments means that measured bacterial abundances will be biased towards organisms that are extracted more efficiently with this method, but this effect should be consistent between conditions in the same experiment. Finally, although we detected DNA mapping to known small plasmids in samples spiked with *K. pneumoniae*, we are unable to resolve the exact plasmid structures or sizes with available sequencing data.

Given the difficulties with the methods tested here, alternative ways of quantifying scarce AMR genes are needed. An alternative to enrichment of organisms is enrichment of the genes themselves, and the approach of targeted capture of DNA fragments has recently been used to increase the abundance of AMR genes up to 300 fold^7^. However, our study demonstrates the need for any method to be validated in a range of well characterised conditions before it can reliably be used to make quantitative comparisons.

### Conclusions

Our study has assessed novel approaches to lowering the limit of detection of AMR genes in *Enterobacteriaceae* in a sequencing-based assay of the intestinal resistome. By comparing different strains of *Enterobacteriaceae* we have demonstrated the problems in using culture enrichment whilst maintaining quantification. Assessing the differential effect of plasmid DNA extraction by plasmid size has highlighted the artefacts that make it difficult to use in a resistome assay requiring accurate quantification. Further attempts at quantitative resistome assessment must either use different methods to amplify scarce AMR genes, or accept the likelihood of missing genes in low abundance organisms, which may include important pathogens such as *Enterobacteriaceae*.

## ACKNOWLEDGMENTS

The research was funded by the National Institute for Health Research Health Protection Research Unit (NIHR HPRU) in Healthcare Associated Infections and Antimicrobial Resistance at the University of Oxford in partnership with Public Health England (PHE) [HPRU-2012-10041], the NIHR Oxford Biomedical Research Centre, and a Medical Research Council UK Clinical Research Training Fellowship to NJF. The views expressed are those of the author(s) and not necessarily those of the NHS, the NIHR, the Department of Health or PHE. DWC and TEAP are NIHR Senior Investigators. Carbapenemase-producing *Enterobacteriaceae* strains provided by Nicole Stoesser. Carbapenemase copy estimation performed by Anna Sheppard.

## Supplementary Figure

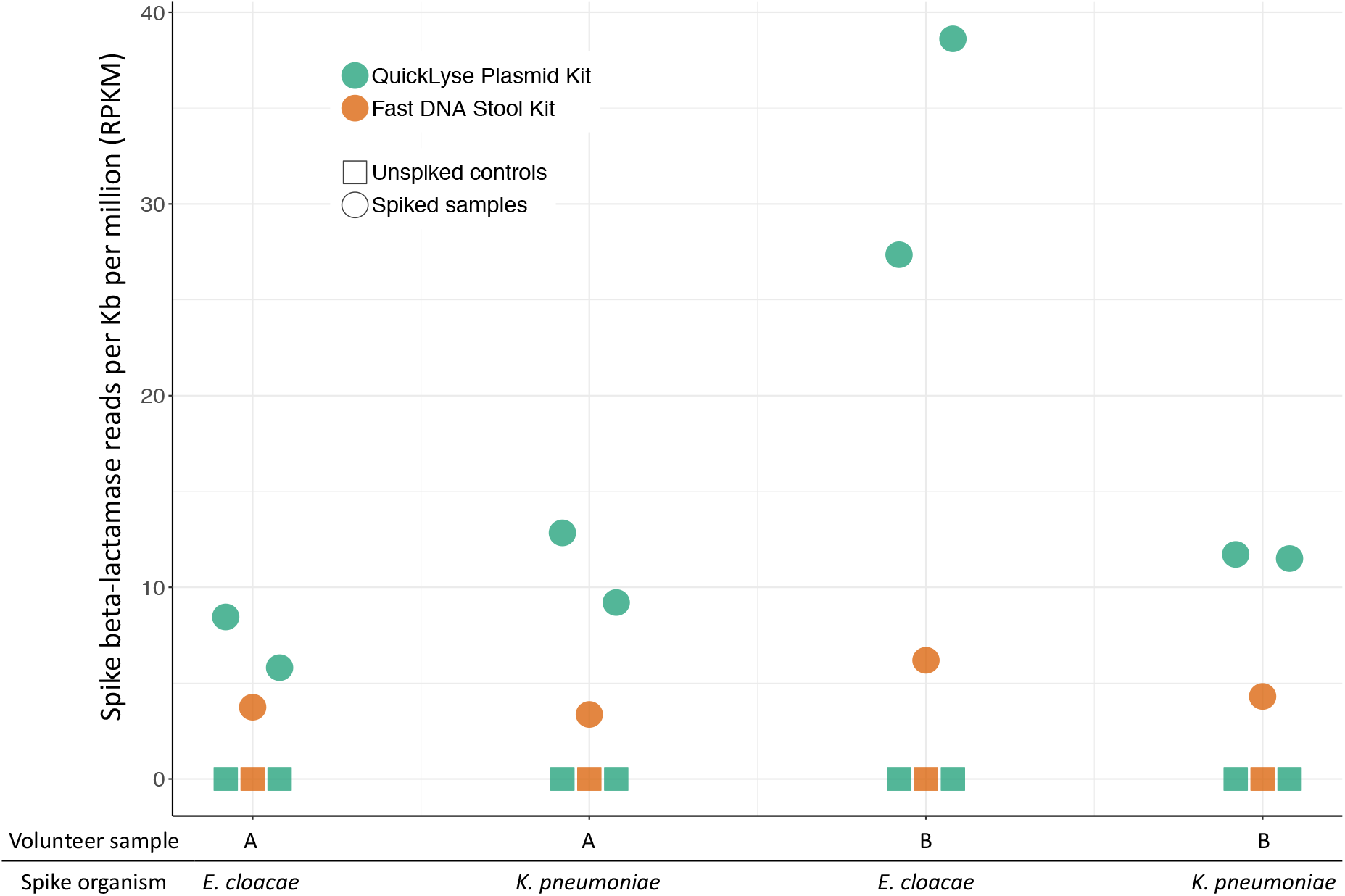
Reads mapping to plasmid-associated beta-lactamases in DNA extracted from faecal samples spiked with 10^8^ CPE/g. Two DNA extraction methods were compared using one of two spike CPE (*K. pneumoniae* & *E. cloacae*) and one of two volunteer samples (A & B). QuickLyse extractions were performed in duplicate.

